# Cis-regulatory architecture of human ESC-derived hypothalamic neuron differentiation aids in variant-to-gene mapping of relevant complex traits

**DOI:** 10.1101/2020.07.06.146951

**Authors:** Matthew C. Pahl, Claudia A. Doege, Kenyaita M. Hodge, Sheridan H. Littleton, Michelle E. Leonard, Sumei Lu, Rick Rausch, James A. Pippin, Jonathan P. Bradfield, Reza K. Hammond, Keith Boehm, Robert I. Berkowitz, Chiara Lasconi, Chun Su, Alessandra Chesi, Matthew E. Johnson, Andrew D. Wells, Benjamin F. Voight, Rudolph L. Leibel, Diana L. Cousminer, Struan F.A. Grant

## Abstract

The hypothalamus regulates metabolic homeostasis by influencing behavior, energy utilization and endocrine systems. Given its role governing health-relevant traits, such as body weight and reproductive timing, understanding the genetic regulation of hypothalamic development and function should yield insights into these traits and diseases. However, given its inaccessibility, studying human hypothalamic gene regulation has proven challenging. To address this gap, we generated a chromatin architecture atlas of an established embryonic stem cell (ESC)-derived hypothalamic-like neuron (HN) model across three stages of *in vitro* differentiation. We profiled accessible chromatin and identified physically interacting contacts between gene promoters and their putative cis-regulatory elements (cREs) to characterize changes in the gene regulatory landscape during hypothalamic differentiation. Next, we integrated these data with GWAS loci for multiple traits and diseases enriched for heritability in these cells, identifying candidate effector genes and cREs impacting transcription factor binding. Our results reveal common target genes for these traits, potentially identifying core hypothalamic developmental pathways. Our atlas will enable future efforts to determine precise mechanisms underlying hypothalamic development with respect to specific disease pathogenesis.

## INTRODUCTION

The hypothalamus is a critical regulator of many physiological functions, including energy homeostasis, reproduction, sleep and stress^1^. This brain region senses neural and physiological signals, which triggers distinct populations of neurons to release neurotransmitters and peptide neuromodulators to signal the autonomic nervous and endocrine systems^1-3^. Monogenic mutations in key nutrient-sensing hypothalamic genes, such as the leptin and melanocortin 4 receptors, result in obesity through dysregulating the neural circuit involved in controlling hunger and satiety, while mutations impacting gonadotrophin-releasing hormone signaling impair the onset of puberty by disrupting pituitary gland signaling^2, 4^.

There is a lack of epigenomic data characterizing the genetic regulatory architecture of the developing and mature human hypothalamus, limiting our ability to translate studies into information directly relevant for disease^5^. Recently, improvements in embryonic and induced pluripotent stem cell differentiation strategies^1, 6, 7^ have partially mitigated the need to study human hypothalamic neurons *ex vivo*. As the precise regulation of hypothalamic development remains poorly understood, differentiating hypothalamic neurons from ESCs provides an opportunity to study these cells and their precursors over time, which should lead to greater understanding of the development of hypothalamic-governed traits and diseases.

Genome wide association studies (GWAS) have yielded hundreds of loci statistically associated with phenotypes known to involve hypothalamic function^8-10^. GWAS efforts typically only report single nucleotide polymorphisms (SNPs) yielding the statistically strongest associations per locus. However, these lead SNPs are not necessarily the causal variants due the presence of other SNPs in linkage disequilibrium (LD). The majority of GWAS signals reside in non-coding regions of the genome, suggesting that their impact on phenotype is primarily via gene regulation. As *cis*-acting regulatory elements, such as enhancers or silencers, can act locally or over large genomic distances, the nearest gene to a GWAS signal may not be the principal effector gene^11-14^. Thus, a major challenge in complex trait genetics is to confidently identify the precise regulatory variant(s) tagged by sentinel SNPs and their corresponding effector target gene(s).

Chromatin conformation approaches to can identify regions harboring SNPs contacting effector genes via long-range promoter interactions in various cell and tissue contexts^16-18^. Recently, we combined a suite of techniques to systematically evaluate GWAS signals located in distal elements^19-22^. Together, our integrated “variant-to-gene mapping” approach aims to physically fine-map significant GWAS loci by identifying open proxy SNPs in LD with each given sentinel signal that directly contact a gene promoter. Assaying relevant cell types in this regard is critical, as promoter architecture varies across cellular identity and developmental-stage^23,16,24^.

While changes in hypothalamic gene expression during development have been studied ^25,26^, the corresponding cis-regulatory architecture in hypothalamus development remains largely unexplored. In this study, we used an arsenal of molecular techniques to characterize the genetic architecture of differentiation of embryonic stem cells, first into hypothalamic progenitors (HPs) and then arcuate (ARC) nucleus-like hypothalamic neurons (HN). We subsequently superimposed GWAS findings for relevant traits on these data to implicate critical and novel effector genes, along with their corresponding putative regulatory elements.

## MATERIALS AND METHODS

### Human ESC-derived hypothalamic neuron differentiation

The HN differentiation protocol was described previously^27^. Briefly, the human ESC H9 line was seeded on Matrigel plates (16 million cells/148 cm^2^; 5 ⨯ 148 cm^2^ Corning dishes) in ESC medium (KnockOut DMEM supplemented with 15% KnockOut Serum Replacement, 0.1mM MEM Non-Essential Amino Acids, 2mM GlutaMAX, 0.06 mM 2-Mercaptoethanol) with FGF-Basic (AA 1 – 155), (20 ng/ml media) and 10 μM Y-27632. Upon confluency (day 1), cells were cultured in ESC medium without FGF-Basic and Y-27632, but supplemented with Shh (100 ng/ml), Purmorphamine (2 μM), 10 μM SB431542, and 2.5 μM LDN193289. From days 5 to 8, ESC medium was gradually replaced with neuroprogenitor medium (DMEM/F-12 supplemented with 0.1 mM MEM Non-Essential Amino Acids, N-2 Supplement, 0.2 μM ascorbic acid, 0.16% glucose). At day 9, cells were switched into neuronal differentiation medium (DMEM/F-12 supplemented with 0.1 mM MEM Non-Essential Amino Acids, N-2 Supplement, B-27 Supplement minus Vitamin A, 0.2 μM ascorbic acid, 0.16% glucose containing 10 μM DAPT). On day 12, cells were collected with TrypLE Express Enzyme at 37°C for 7 min and washed twice including filtration through pre-wetted 40 μm Corning Sterile Cell Strainer. The hypothalamic progenitor cell pellet was then resuspended with neuronal differentiation medium containing 10 μM Y-27632 for plating on 148 cm^2^ dishes coated with Poly-L-Ornithine solution (0.01%) and laminin (4 μg/ml) at a seeding density of 16 million cells/148 cm^2^. After 4 hours, the medium was changed to neuronal differentiation medium supplemented with 10 μM DAPT. On day 15, the neuronal differentiation medium was supplemented with 20 ng/ml BDNF until collection on day 27.

### ATAC-seq, RNA-seq, Capture C Library generation, Processing, Peak Calling ATAC-seq library generation, analyses, and peak calling

#### ATAC-seq library generation

A total of 50,000 cells were centrifuged at 550g for 5 min at 4°C. The cell pellet was washed with cold PBS and resuspended in 50 μL cold lysis buffer (10 mM Tris-HCl, pH 7.4, 10 mM NaCl, 3 mM MgCl2, 0.1% NP-40/IGEPAL CA-630) and immediately centrifuged at 550g for 10 min at 4°C. Nuclei were resuspended in the Nextera transposition reaction mix (25 μL 2x TD Buffer, 2.5 μL Nextera Tn5 transposase, and 22.5 μL nuclease free H2O) on ice, then incubated for 45 min at 37°C. The tagmented DNA was then purified using the Qiagen MinElute kit and eluted in 10.5 μL Elution Buffer (EB). Ten microliters of purified tagmented DNA was PCR amplified using the Nextera Indexing Kit for 12 cycles to generate each library. The PCR reaction was subsequently purified using 1.8x AMPure XP beads, and concentrations were measured by Qubit Fluorometer. Quality of completed libraries was assessed on a Bioanalyzer 2100 high sensitivity DNA Chip. Libraries were paired-end sequenced at the Center for Spatial and Functional Genomics on the Novaseq 6000 platform (50 bp read length).

#### Analysis and Peak calling

The number reads from the hypothalamic neurons were down-sampled to make the sequencing depth comparable between conditions using sambamba. Open chromatin regions were called using the ENCODE ATAC-seq pipeline. Pair-end reads from all replicates for each cell type were aligned to the hg19 genome using bowtie2, and duplicate reads were removed from the alignment. Aligned tags were generated by modifying the reads alignment by offsetting +4bp for all the reads aligned to the forward strand, and -5bp for all the reads aligned to the reverse strand. Narrow peaks were called independently for pooled replicates for each cell type using macs2 (-p 0.01 --nomodel --shift -75 --extsize 150 -B --SPMR --keep-dup all --call-summits) and ENCODE blacklist regions were removed from called peaks. We then merged peaks with at least 1 bp overlap between replicates to generate a consensus set of peaks. The consensus set peaks were filtered to those which were reproducible in at least half the ATAC-seq replicates using bedtools intersect^11^. For analyses involving cell-type specific sets of peaks, we considered the set of consensus peaks with mean FPKM value greater than 1 to be “open” in that cell type.

For TF analysis replicated, deduplicated ATAC-seq bam files were merge and downsampled to consistent read count for each stage of differentiation to calculate purity scores for each TF.

#### Differential analysis of chromatin accessibility

To identify differentially accessible OCRs between ESCs, HPs, and HNs, we used the R package csaw, which uses the de-duplicated read counts for the consensus OCRs for each replicate to normalized against background (10K bins of genome). OCRs with median value of less than 1.2 CPM (∼10-50 reads per OCR) across all replicates were removed from further differential analysis. Similar to RNA-seq differential analysis, accessibility differential analysis of the consensus OCRs was performed using glmQLFit approach, fitting cell type in edgeR, but using the csaw normalization scaling factors. Differential OCRs between cell types were identified with thresholds of FDR < 0.05 and absolute log2 fold change > 1. FPKM values were calculated for all OCRs in the consensus list.

### RNA-seq library generation and analysis

#### RNA-seq library generation

RNA was isolated from each cell type in triplicate using TRIzol Reagent. RNA was then purified using the Direct-zol RNA Miniprep Kit and depleted of contaminating genomic DNA using DNAse I. Purified RNA was then checked for quality on the Bioanalyzer 2100 using the Nano RNA Chip and samples with a RIN number above 7 were used for RNA-seq library synthesis. RNA samples were depleted of rRNA using the QIAseq FastSelect RNA Removal Kit then processed into libraries using the NEBNext Ultra II Directional RNA Library Prep Kit for Illumina according to manufacturer’s instructions. Quality and quantity of the libraries was measured using the Bioanalyzer 2100 DNA chip and Qubit Fluorometer. Completed libraries were pooled and sequenced on the NovaSeq 6000 platform using paired-end 51bp reads at the Center for Spatial and Functional Genomics at CHOP.

#### RNA-seq processing and differential expression analysis

Sequencing data was demultiplexed and FastQ files were generated using Illumina bcl2fastq2 conversion. Paired-end Fastq files with for each replicate were mapped to the reference genome using STAR. Gene features were assigned to a curated annotation consisting of GencodeV19 with lincRNA and sno/miRNA annotation from the UCSC Table Browser. The raw read count for each gene feature was calculated using HTSeq-count. with parameter settings -f bam -r pos -s reverse -t exon -m intersect. The genes located on chrM or annotated as ribosomal RNAs were removed before further processing.

Differential analysis was performed in R using the edgeR package. Briefly, the raw reads per genes features were converted to read Counts Per Million mapped reads (CPM). The gene features with median value of less than 0.7 CPM (10 ∼ 18 reads per gene feature) across all samples were filtered. Normalization scaling factors were calculated using the trimmed mean of M-values method. Differentially expressed genes between ESCs, HPs, and HNs were identified with thresholds of FDR < 0.05 and absolute log_2_FC > 1. Expression values are reported as transcript per million mapped reads (TPM). We clustered standardized TPM values of differentially expressed genes using the R function hclust. Genes expression values were standardized using the R function, scale. Following this, the top six branches were cut to define the clusters used in subsequent comparisons.

### Capture C Library Generation and Analysis

#### 3C Library Generation

We used standard methods for 3C library generation^82^. For each library, 10^7^ fixed cells were thawed at 37°C, followed by centrifugation at RT for 5 mins at 14,000 rpm. The cell pellet was resuspended in 1 mL of dH2O supplemented with 5 μL 200X protease inhibitor cocktail, incubated on ice for 10 mins, then centrifuged. Cell pellet was resuspended to a total volume of 650 μL in dH2O. 50 μL of cell suspension was set aside for pre-digestion QC, and the remaining sample was divided into 3 tubes. Both pre-digestion controls and samples underwent a pre-digestion incubation with the addition of 0.3% SDS, 1x NEBuffer DpnII, and dH2O for 1hr at 37°C in a Thermomixer shaking at 1,000rpm. A 1.7% solution of Triton X-100 was added to each tube and shaking was continued for another hour. After the pre-digestion incubation, 10 μL of DpnII was added only to each sample tube, and continued shaking along with the pre-digestion control until the end of the day. An additional 10 µL of DpnII was added to each digestion reaction and digestion continued overnight. The next day, another additional 10 µL of DpnII was added and the incubation continued for another 2-3 hours. 100 μL of each digestion reaction was then removed, pooled into one 1.5 mL tube, and set aside for digestion efficiency QC. The remaining samples were heat inactivated at 65°C for 20 min at 1000 rpm in a Thermomixer, and cooled on ice for 20 additional minutes. Digested samples were ligated with 8 μL of T4 DNA ligase and 1X ligase buffer at 1,000 rpm overnight at 16°C in a Thermomixer. The next day, an additional 2 µL of T4 DNA ligase was spiked into each sample and incubated for another few hours. The ligated samples were then de-crosslinked overnight at 65°C with Proteinase K along with the pre-digestion and digestion controls. The following morning, both controls and ligated samples were incubated for 30 min at 37°C with RNase A, followed by phenol/chloroform/isoamyl alcohol (Fisher Cat # BP1752I400) extraction and ethanol precipitation at -20°C. The 3C libraries were centrifuged at 3000 rpm for 45 min at 4°C, while the scontrols were centrifuged at 14,000 rpm, to pellet the samples. DNA pellets were resuspended in 70% ethanol and again centrifuged as described above. The 3C library pellets and control pellets were resuspended in 300 μL and 20 μL dH2O, respectively, and stored at −20°C. Sample concentrations were measured by Qubit Fluorometer. Digestion and ligation efficiencies were assessed by gel electrophoresis on a 0.9% agarose gel and quantitative PCR (Brilliant III SYBR qPCR Master Mix, VWR Cat # 97066-528).

#### Promoter Capture Library Generation

We followed our same protocols as previously published^23^. Isolated DNA from 3C libraries was quantified using a Qubit Fluorometer, and 10 μg of each library was sheared in dH2O using a QSonica Q800R to an average fragment size of 350bp. QSonica settings used were 60% amplitude, 30s on, 30s off, 2 min intervals, for a total of 5 intervals at 4 °C. After shearing, DNA was purified using AMPure XP beads. DNA size was assessed on a Bioanalyzer 2100 using a DNA 1000 Chip and DNA concentration was checked via Qubit Fluorometer. SureSelect XT library prep kits were used to repair DNA ends and for adaptor ligation following the manufacturer protocol. Excess adaptors were removed using AMPure XP beads. Size and concentration were checked again by Bioanalyzer 2100 using a DNA 1000 Chip and by Qubit Fluorometer before hybridization. One microgram of adaptor-ligated library was used as input for the SureSelect XT capture kit using manufacturer protocol and our custom-designed 41K promoter Capture-C probe set. The quantity and quality of the captured libraries were assessed by Bioanalyzer using a high sensitivity DNA Chip and by Qubit Fluorometer. SureSelect XT libraries were then paired-end sequenced on Illumina NovaSeq 6000 platform (51bp read length) at the Center for Spatial and Functional Genomics at CHOP.

#### Analysis of Capture C

Paired-end reads from three replicates from ESCs, HPs, and HNs were pre-processed using the HICUP pipeline with the default parameters. Reads were aligned to hg19 using bowtie2. We called call significant promoter interactions using the read count from promoters included in our reference bait. As previously reported^23^, significant interactions were called using CHiCAGO with default parameters except for bin-size set to 2,500. In addition to our analysis per individual DpnII fragment (1frag), we also called interactions by binning four fragments, which improves detection of long-distance interactions. Significant interactions at 4-DpnII fragment resolution were also called using CHiCAGO. Interactions with a CHiCAGO score > 5 in at least one cell type in either 1-fragment or 4-fragment resolution were considered significant.

Quality control metrics: Reproducibility between ATAC-seq and RNA-seq samples was determined by principal component analysis and pairwise pearson correlation coefficients between samples. Median expression values were downloaded from GTeX (v7). The spearman rank correlation of the genes with expression level pass a threshold of TPM > 5 in at least one cell/tissue type (16,953 genes) were calculated using Spearman’s Correlation Coefficient (cor function in R).

Genome tracks were visualized using the python packager pyGenomeTracks version 3.0. Genomic Annotations: Promoters were defined as 1500kb upstream and 500kb downstream of the TSS (Genecode V19). Overlapping annotations were assigned to genomic features based on a hierarchy of (1) Promoter, (2) 5’UTR, (3) CDS, (4) 3’UTR, (5) first intron, (6) other introns, or (7) intergenic. The percentage of OCRs overlapping with each feature was visualized as pie charts using ggplot2.

#### Genomic coordinate visualization

All coordinates referring to hg19 as the reference genome.

### Variant to gene mapping

We identified proxies located in open chromatin and fragments interacting with a bait using bedtools intersect, as performed in our previously published method^18^. We considered all interactions with a proxy SNP located in a distal interaction fragment and those falling within OCRs located in baits. Putative target effector genes were then filtered by expression in each respective cell state (TPM > 1). These genes were functionally annotated by the DAVID functional annotation tool.

### Gene set enrichment

GO and REACTOME datasets annotated in MSigDB (v7.0) were used for gene set enrichment analyses. Statistical significance of gene set enrichment was determined using the hypergeometric test, implemented in the R phyper function.

### Bioinformatic Analyses

Transcription factor analysis: PIQ, which integrates TF motif scanning with TF footprinting using DNAase or ATAC-seq data, was used to predict TF binding sites^28^. We scanned JASPAR2020 core^29^ PWMs against hg19, with ENCODE blacklist regions excluded using the default settings. For downstream analyses we considered TF binding sites passing the default cutoff of purity > 0.7. We identified TF motifs enriched in cREs compared the set of nonPIR-OCRs using the R package BiFET, with a cutoff of FDR<0.05,

#### Partitioned LD Score Regression

Partitioned heritability was measured using LD Score Regression v1.0.0^30^. Partitioned LDSR requires the GWAS summary statistics and a feature annotation. ESC, HP, and HN annotations were generated using bed files containing positions of the cRE (promoter OCRs + PIR-OCRs) with +/-500bp extension as previously performed^30^.

We selected a set of traits related to metabolic, endocrine, and neuropsychiatric traits with available GWAS summary statistics **(see Supplemental methods)**.

#### Variant to gene mapping

Sentinel SNPs were collected from the most recent large-scale GWAS studies. Proxies for each sentinel were queried using SNiPa using the following parameters: Genome assembly GRCh37; Variant set 1000 Genomes, Phase 3 v5; Population European; Genome annotation Ensembl 87 and r^2^ > 0.6. Intersection as done previously (see Supplemental Methods). Analysis was then restricted to expressed genes with a TPM > 1.

#### GWAS Colocalization

Summary statistics for 6 regions with overlapping associations for 3-4 input traits were imputed using FIZI. Common variants (MAF ≥ 0.01) from the European ancestry 1000 Genomes Project v3 samples were used as a reference panel for the imputation. Default parameters were used with the exception that the minimum proportion parameter was lowered to 0.01. Standard errors and betas for the imputed SNPs were estimated using the method from https://github.com/zkutalik/ssimp_software/blob/master/extra/transform_z_to_b.R. Subsequently, HyPrColoc was used to test for colocalization across all input traits simultaneously. Separately, we tested for colocalization for each input trait genome-wide against GTEx v.7 hypothalamic eQTLs using coloc^31^.

## RESULTS

### ESC-derived hypothalamic-like neurons (HN) recapitulate molecular characteristics of the hypothalamus

We utilized an established protocol to derive ARC HN-like neurons from a human ESC line (H9)^32, 33^, and collected cells at three stages of differentiation: pluripotent ESCs, NKX2-1+ hypothalamic progenitors (HPs), and HNs. We then profiled global gene expression patterns for these three stages using RNA-seq, chromatin accessibility with ATAC-seq, and chromatin conformation via promoter-focused Capture C to generate a high-resolution atlas of the distal promoter interaction landscape in an *in vitro* human model of hypothalamic development (**Figure 1A**). To assess the reproducibility between replicates, we performed principal component analysis and pairwise Pearson correlation on the RNA-seq and ATAC-seq datasets. In both cases the first principal component corresponded to the stage of differentiation and accounted for more than half of the variation (RNA-seq: 52.60%; ATAC-seq: 55.30%) (**Supplemental Figure 1A-D**).

**Figure 1:**
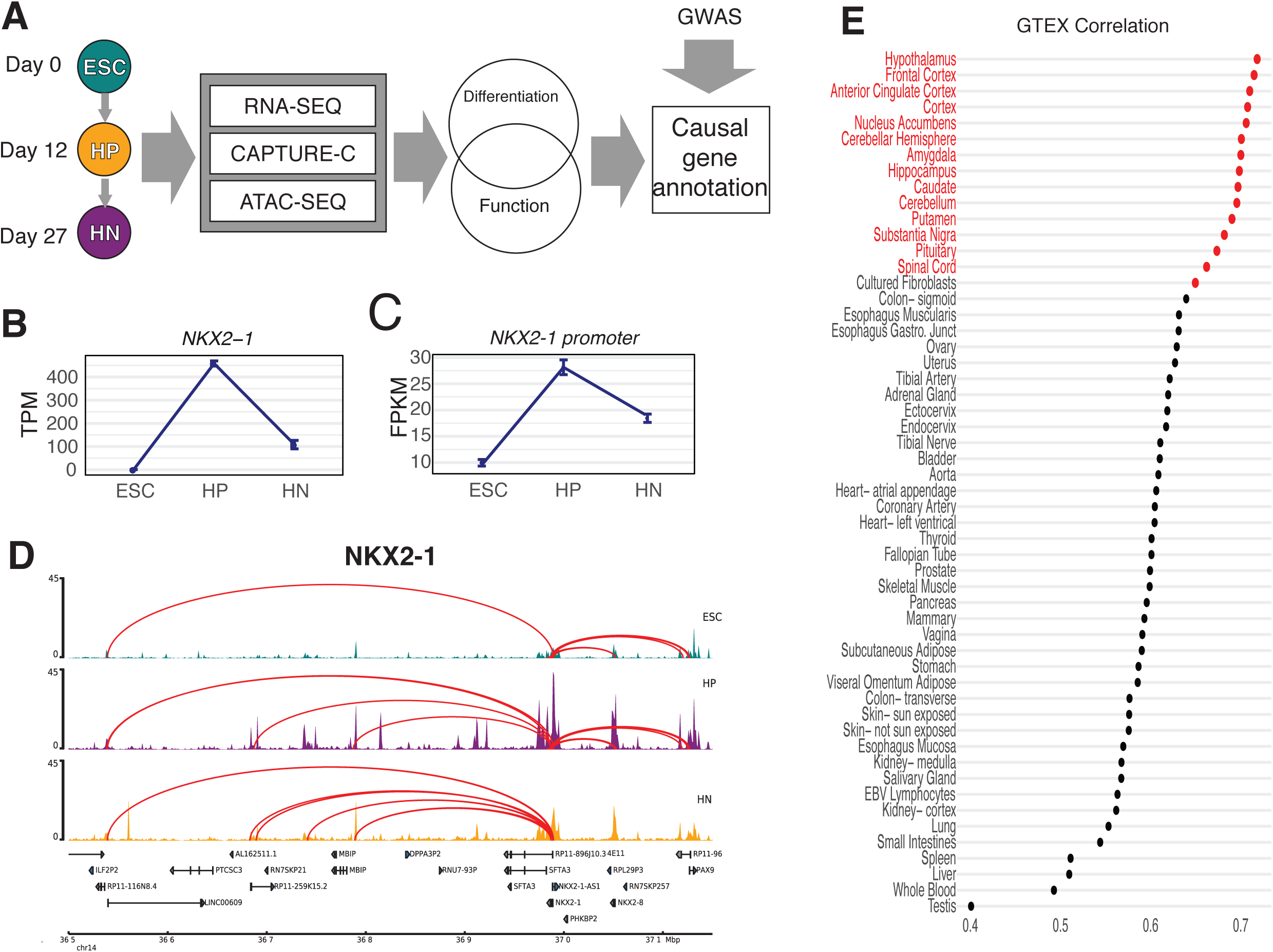
An integrative functional genomics approach to model the differentiation of hypothalamic neurons. (A) Schematic of the study design. ESC, HP, and HNs were used to generate RNA-seq, ATAC-seq, and Capture C profiles, which we compared to the GWAS signals mined using our variant to gene mapping approach. (B) Expression level of NKX2-1 determined by RNA-seq. (C) Accessibility change of OCR located in the NKX2-1 promoter over the course of HN differentiation. (D) A 600kb region around the NKX2-1 gene in ESCs (teal), HPs (purple), and HNs (orange). The peaks track represents the ATAC-seq coverage, where higher peaks depict increased accessibility (open chromatin), and arcs represent significant (Chicago Score > 5) interacting regions between the NKX2-1 promoter. (E) Comparison of HN expression profile to median GTeX database, scores give the spearman correlation coefficient of the top 16,953 genes expressed in both datasets.

To further confirm the molecular congruence of HN differentiation to the *in vivo* development of HNs, we examined the expression of several marker genes (**Figure 1B; Supplemental Figure 1E**)^25^, which were consistent with expectations ^27, 32, 33^. In particular, we investigated the expression and promoter interaction landscape of *NKX2-1*, which encodes a transcription factor (TF) critical for hypothalamus specification. *NKX2-1* is expressed in the developing hypothalamus and subsequently becomes restricted to a subset of neurons^34^. *NKX2-1* expression followed a similar pattern during HN differentiation (**Figure 1B**). In addition, we observed a distinct change in the accessibility of the *NKX2-1* promoter, concordant with its expression pattern (**Figure 1C**), as well as fewer interactions detected as *NKX2-1* expression decreased (**Figure 1D**), confirming our detection of expected dynamic changes.

In addition to confirming the expression of known marker genes, we compared the global transcriptomic profile of HNs to the GTeX RNA-seq database, which is derived from primary human tissue samples (**Figure 1E; Supplemental Table 1**)^35^. HN gene expression was highly correlated with the hypothalamus (Spearman’s rho= 0.719; adjusted *P* = 1.14 × 10^−14^). Taken together, these results show that ESC-derived HNs resemble hypothalamus tissue.

### Temporal dynamics of regulation of gene expression and cis-regulatory elements during hypothalamic neuron differentiation

We assessed the temporal profile of gene expression to identify genes with developmental stage-restricted expression during human HN differentiation, with 15,808 genes differentially expressed in at least one stage **(Figure 2A)**. We assigned these genes to six clusters based on expression patterns during the course of differentiation. Each cluster corresponded to genes specifically enriched or depleted in at least one stage of differentiation (**Figure 2B; Supplemental Figure 2A-B**). Gene Ontology (GO) and REACTOME enrichment analysis of each cluster identified known biological processes related to proliferating progenitor cells and differentiated neurons^36^ (**Supplemental Figure 2 C-D**).

**Figure 2:**
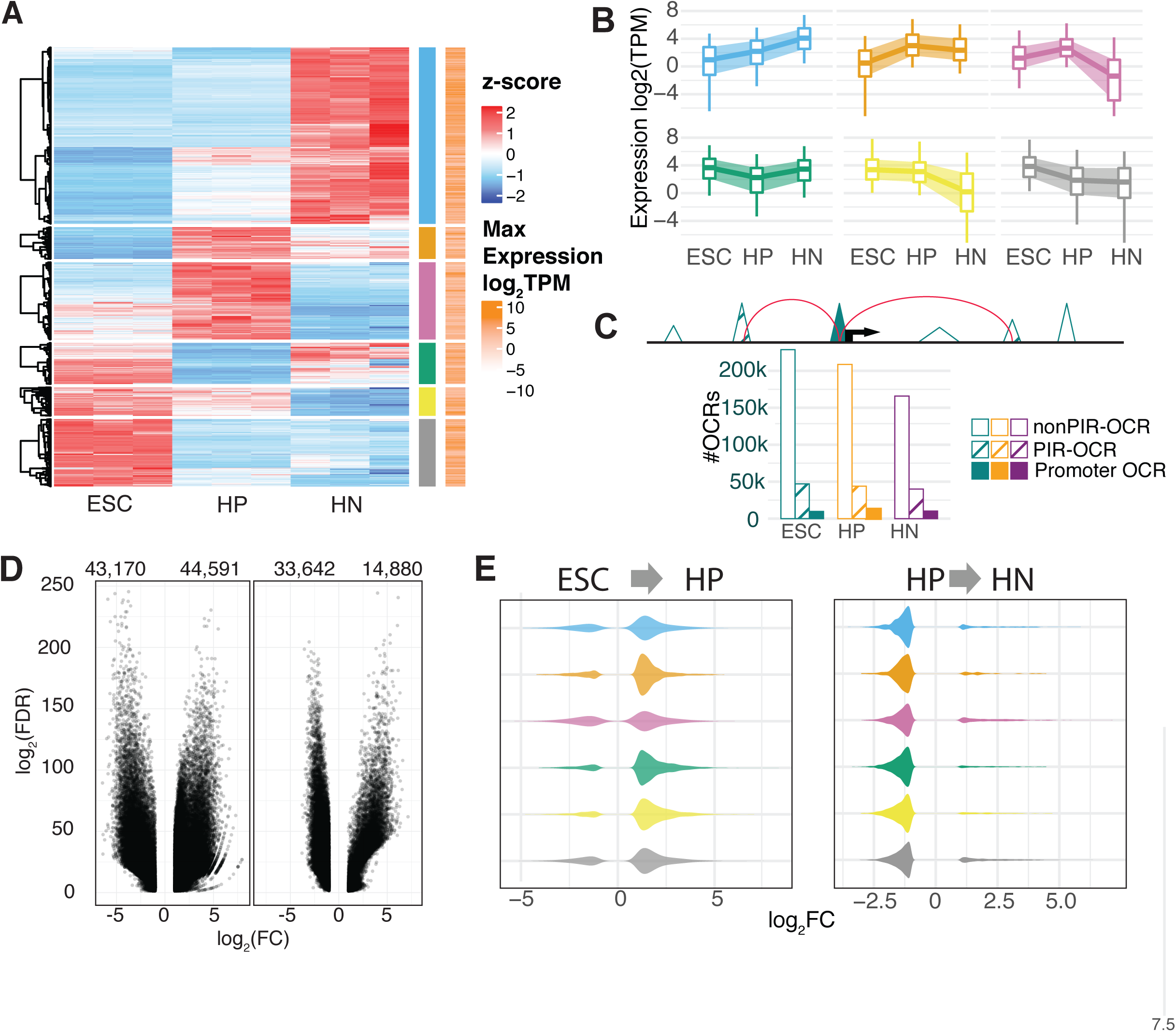
Gene expression and chromatin architecture underlies during hypothalamic neuron differentiation: (A) Heatmap of the scaled expression values (z-score) of each RNA-seq sample of ESC, HP, and HN for the 15,808 genes with differential expression in at least one stage of differentiation. Red indicates higher relative expression, while blue indicates lower relative expression. From hierarchical clustering, the genes were assigned to six groups based on their expression pattern pointed to being either specifically enriched or depleted in one condition (blue, orange, pink, green, yellow, gray bars). The log transformed condition-wise max TPM value for each gene is plotted where white indicates lower expression and orange indicates higher expression compared to other genes. (B) The global average of expression values (TPM) the genes in each cluster. The central line of each boxplot represents the median, with edges representing the 25 and 75 percentiles, and whiskers represent the 5 and 95 percentiles. (C) Sets of OCRs were assigned based on location: promoter OCRs (solid), PIR-OCRs (hashed), and nonPIR-OCRs the remaining OCRs that were not annotated to a gene. Bottom: The distribution of number OCRs annotated to each set per celltype. (D) Volcano plot depicting the global genome-wide significant differentially accessible OCRs in the transition from ESC to HP (left) and HP to HN (right). (H) Distribution of chromatin accessibility fold-change of cisRE annotated to DE gene clusters (**Figure 2A**).

To correlate gene expression changes during HN differentiation with the respective chromatin accessibility and conformation profiles at each stage, we defined the relationship with open chromatin regions (OCRs) using ATAC-seq. We identified a total of 404,691 OCRs in at least one stage. The OCRs were disproportionately located in promoters (−1500/+500bp TSS) and first introns **(Supplemental Figure 3A)**, which is characteristic of regulatory elements^37^. We then grouped the OCRs into three categories **(Figure 2C)**: (1) OCRs located within promoter regions annotated as ‘promoter OCRs’; (2) OCRs with direct promoter contacts determined by Capture C, annotated as ‘promoter-interacting region (PIR)-OCRs’; and (3) OCRs that could not be assigned to a gene because they did not fit either criteria, annotated as ‘nonPIR-OCRs’. Because they could be annotated to a gene, we considered the sets of 50,952 promoter OCRs and 87,170 PIR-OCRs as putative cREs.

Both the number of cREs per gene and the mean distance between the cRE and the promoter were decreased in HPs compared to ESCs or HNs **(Supplemental Figure 3B-C)**, reflecting fewer long-range interactions detected at this stage **(Supplemental Figure 3D-E)**. We also observed a trend for genes with higher expression interacting with more cREs (Kruskal Wallis test: p-value < 2.2 x10^−16^) **(Supplemental Figure 3F)**, which is in line with reports for other neuronal^38^ and immune cells^22^.

We then compared chromatin accessibility across the three stages, and identified 87,761 differentially accessible regions from ESCs to HPs (43,170 more closed; 44,591 more open) and 48,522 differentially accessible regions from HPs to HNs (33,642 more closed; 14,880 more open) **(Figure 2G; Supplemental Figure 3G)**. The genome-wide decrease in open chromatin as cells advanced in differentiation occurs in other developing tissues^39^. The subset of cREs contacting differentially expressed genes showed a net increase in accessibility as ESCs differentiated into HPs, and subsequently decreased as HPs advanced to HNs, regardless of gene expression pattern defined by our clustering analysis (**Figure 2H; Figure 2A**). This trend suggests that regulatory elements driving the gene expression changes specifying undifferentiated progenitors to a hypothalamic cell fate are primarily first established by opening of selected cREs followed by a more global pruning of contacts upon differentiation to neurons.

### Predicting transcription factors controlling HN development from spatial gene regulatory architecture

TFs regulate gene expression by binding to specific DNA sequences such as enhancers and silencers. Local chromatin accessibility is a critical determinant of where and when TFs bind to DNA^40^. To identify TFs that may bind to cREs, we leveraged PIQ, which uses chromatin accessibility profiles to improve motif score-based matching ^28^. We identified putative binding sites in each stage of differentiation and observed that more binding sites were detected in HNs compared to ESCs or HPs (**Figure 3A)**. After grouping TFs by family, we detected more binding sites for Homeodomain TF factors in HNs compared to ESCs or HPs (**Figure 3B)**. This result was expected, as neuronal identity is refined by the expression of multiple patterning genes^41^. We also observed fewer AP-1 family binding sites in HNs compared to ESCs and HPs; these TFs regulate the cell cycle in early cellular development^42^.

**Figure 3:**
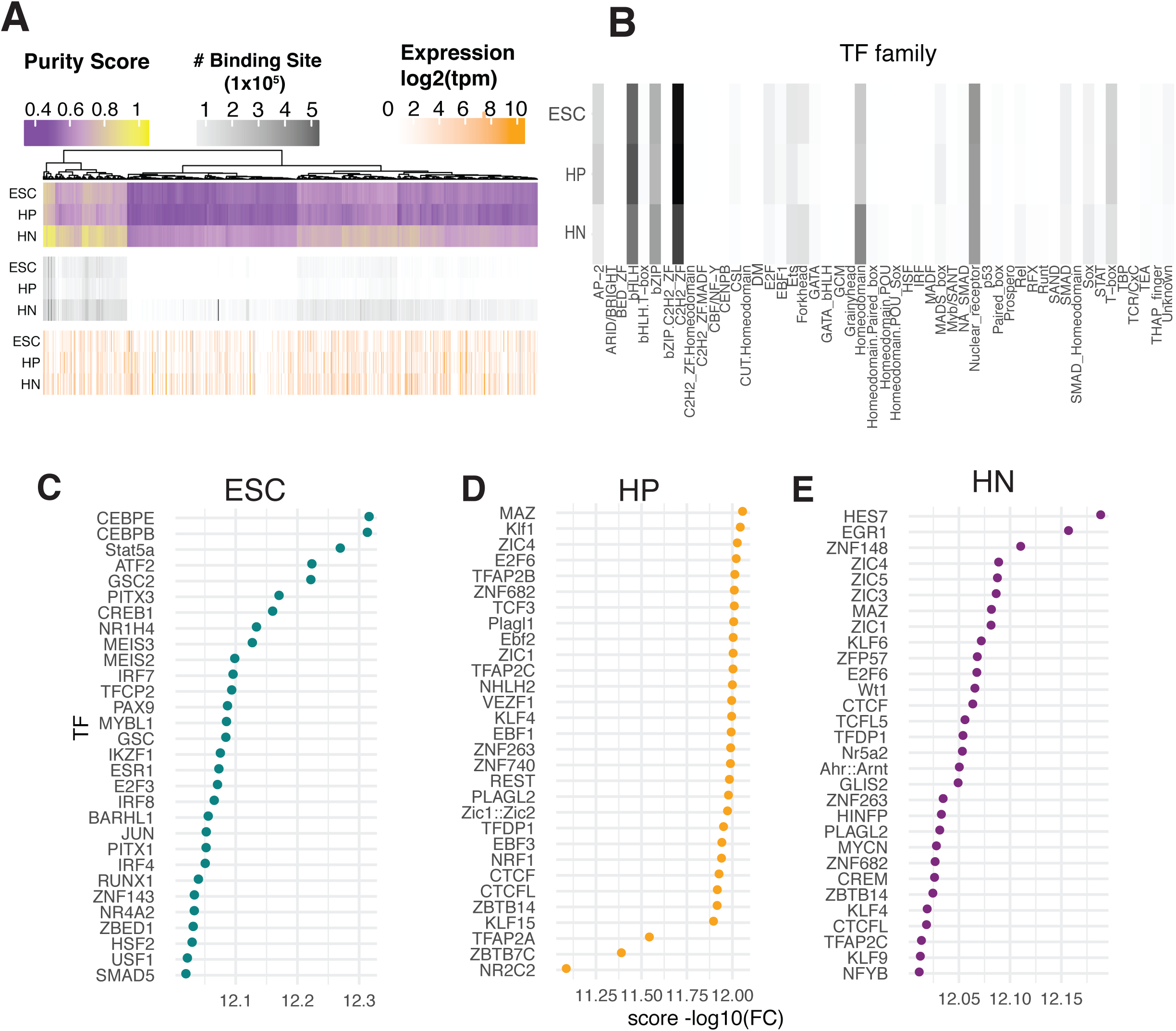
TF footprinting and analysis: **(A)** The summary of TF binding site prediction, the Purity scores calculated by PIQ (Purple/Yellow), the number of TF binding sites passing purity cutoff > 0.7 threshold (Grayscale), log_2_ transformed expression of the respective TF in HN (white /Orange). (B) Grouping number of TF binding sites family. Darker color indicates a higher number of predicted binding sites. (C) Enrichment of TF binding sites in putative cREs compared to other OCRs in each cell type adjusted for GC content and read count.

Next, we checked for TF enrichment in cREs in each cell type to identify which TFs could mediate these promoter contacts (**Figure 2C**). We compared the three stages of differentiation for enriched binding in the cREs compared to non-PIR OCRs. This approach generated a set of potentially relevant TFs involved in HN differentiation. We found 474 enriched TFs in ESCs, 122 in HPs and 134 in HNs (**Figure 3C-E; Supplemental Table 2**). While some TFs involved in DNA looping, such as MAZ and CTCF, were enriched in all three cell types, we also observed differences in the top enriched TFs in each comparison such as ZBTB6, EBF2, EGR1, and ZIC/MYC family members (**Supplemental Figure 4**).

### GWAS loci enriched in hypothalamus cREs

We hypothesized that cREs in our hypothalamic model are enriched for genetic variants associated with traits that are at least partly governed by the hypothalamus. We used Partitioned Linkage Disequilibrium Score Regression (LDSR)^30^ to identify significantly enriched traits for associated loci falling into hypothalamic cREs. We assembled GWAS summary statistics from several recent studies examining metabolic, circadian, neuropsychiatric and puberty-relevant phenotypes, and tested whether cREs were enriched for GWAS loci in at least one stage of differentiation (**Figure 4A**). We detected significantly enriched signals with BMI, adult height, age at menarche (AAM), major depressive disorder (MDD), bipolar disorder, several measures of sleep (**Figure 4A; Supplemental Table 3**). The enrichment of GWAS loci for these hypothalamus-related traits in the cREs identified in the ESC-derived hypothalamic cell types highlights their utility as a model for gaining insight into the target effector genes and regulatory elements functionally related to these traits and diseases.

**Figure 4:**
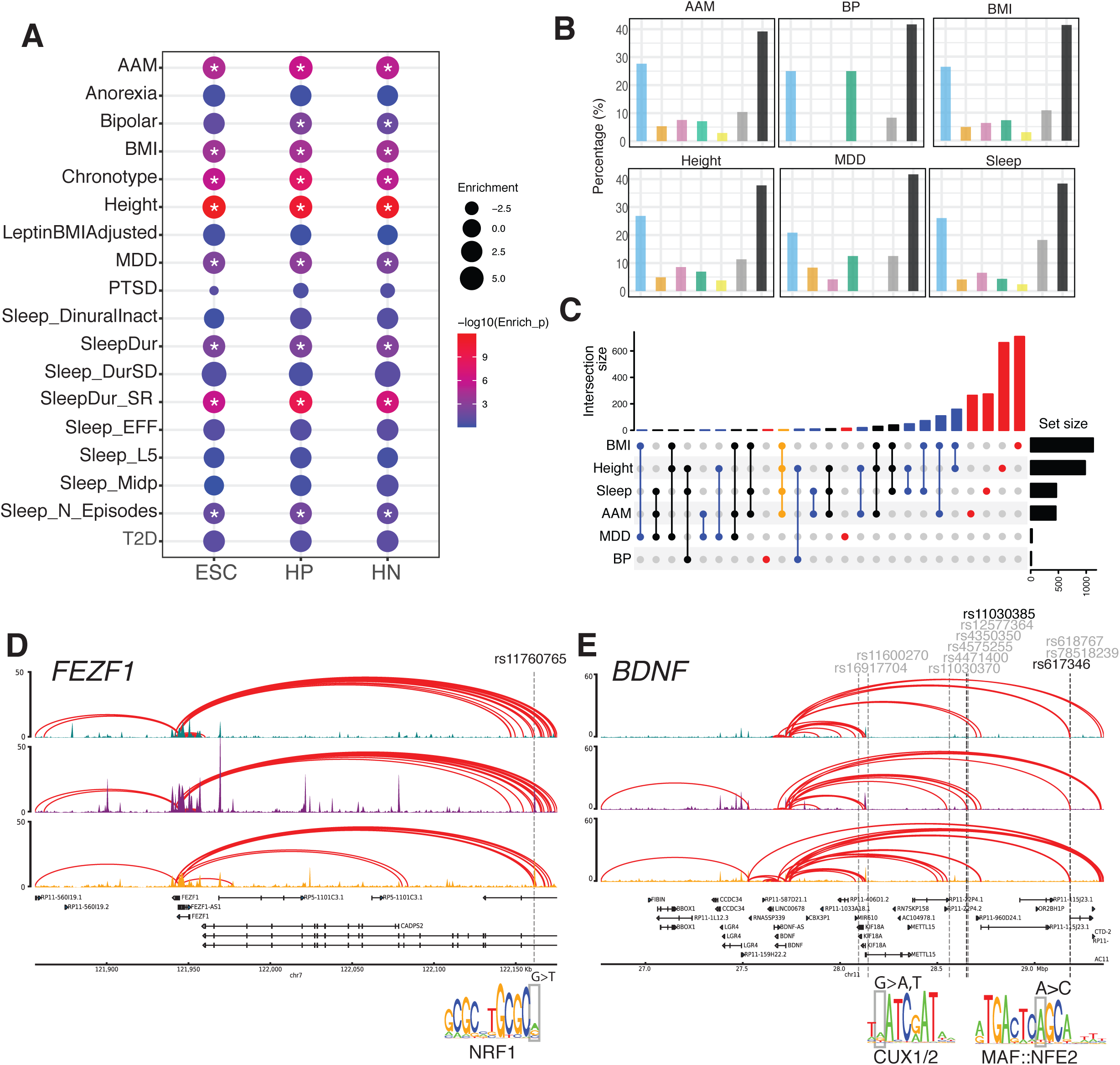
Putative cis regulatory elements are enriched for GWAS signals of complex traits. (A) Partitioned LDscore regression of ESC, HP, and HN cRE against the indicated genome-wide signal. Circle size indicates the enrichment of estimated heritability and color indicates statistical significance. (B) Comparison of genes implicated in our variant to gene mapping analysis for each GWAS. Dots and lines indicate the intersect of the set of genes found in each GWAS. Top: the number of genes in different overlapping sets. Right: the number of SNPs detected in each GWAS. (C) Proportion of genes from our variant to gene mapping located in each DE cluster (**Figure 2A**) or non-DE genes (black). (D) *FEZF1* genomic locus with interactions connecting to a distal cRE. The SNP is located in a putative NRF1 motif. (F) Genome track for the BDNF locus. Multiple proxies in open regions are shown. Two proxies located in putative TF motifs for CUX1/2 and MAF::NFE2 are shown. Abbreviations: AAM: Age at Menarche; BMI: Body Mass Index; LeptinBMIadj: Circulating Leptin levels adjusted for BMI; MDD: Major Depressive Disorder; PTSD: Post Traumatic Stress Disorder; Sleep L5 Time; Least active 5 hours; Sleep N Episodes = Number of Sleep episodes; Sleep Dur; Sleep Duration; Sleep Dur SD: Sleep Duration Standard Deviation; Sleep Diurnal Inact: Sleep Diurnal Inactivity; T2D: Type 2 Diabetes.

### Variant-to-gene mapping identifies target effector genes at GWAS-implicated loci

Guided by the results of the partitioned LDSR analyses, we performed variant-to-gene mapping for those traits that displayed significant heritability enrichment in at least one of the three cellular differentiation stages. We began with all genome-wide significant loci in the most recent large-scale GWAS for each respective trait and queried for proxy SNPs in LD with each sentinel SNP. We overlapped this set of SNPs with the open chromatin regions identified by ATAC-seq, and queried our promoter focused Capture C data to determine the genes in physical contact with open proxy SNPs in each of the three cell states. Finally, we filtered by expression from our RNA-seq data to limit subsequent analyses to genes expressed in at least one stage of differentiation (TPM > 1) (**Table 1**).

| Trait | GWAS Signals | In reference panel | Unique proxies | Open proxies with *cis* interactions<sup>a</sup> | Sentinels with *cis* interactions<sup>a</sup> | Contacted genes<sup>b</sup> | Unique contacted genes<sup>b</sup> |
| --- | --- | --- | --- | --- | --- | --- | --- |
| Height | 3290 | 3254 | 165039 | 1081, 1098, 1073 | 594, 562, 564 | 543, 439, 507 | 981 |
| BMI | 941 | 928 | 63148 | 619, 474, 575 | 271, 211, 248 | 800, 526, 802 | 1126 |
| AAM | 499 | 463 | 13040 | 242, 194, 258 | 109, 105, 114 | 365, 267, 268 | 452 |
| Sleep | 459 | 445 | 26445 | 313, 223, 305 | 98, 84, 99 | 361, 212, 319 | 461 |
| MDD | 44 | 42 | 3059 | 22, 18, 17 | 11, 10, 8 | 15, 16, 16 | 24 |
| BP | 39 | 30 | 1697 | 1, 1, 1 | 1, 1, 1 | 2, 0, 11 | 12 |

<sup>a</sup>ESCs, HPs, and HNs, respectively
<sup>b</sup>Expressed at TPM > 1

For each trait, we noticed that multiple contacted genes also have previously characterized relevant monogenic disease mutations, suggesting that our approach can identify genes with known mechanistic links to the queried traits (**Supplemental Table 4**). For BMI, we detected genes that are known to influence monogenic forms of extreme body weight, including *ABCC8*^44^, *BDNF*^*45*^, and *PPARG*^46^. From a AAM locus, we observed *FEZF1*, known to harbor monogenic mutations that cause delayed/absent puberty^47^. Finally, among sleep traits, our data implicated *PER2*, which encodes a factor that plays a role in advanced sleep phase syndrome^48^. Many additional putative effector genes also have plausible biological links to each trait, while others represent novel findings in the context of these phenotypes (**Supplemental Table 5**).

### Variant contacted genes in hypothalamic development

To identify variants impacting hypothalamus development, we compared our contacted SNPs with the set of cREs connected to specific genes for each trait, and found that ≥50% of the SNP contacted genes were differentially expressed during HN differentiation (**Figure 4B**). To identify biological functions associated with these genes, we tested for GO term enrichment specific to either HP or HNs. HPs were enriched for ERK1/ERK2 cascade and phospho-inositol 3 lipid signaling (**Supplemental Table 6**), which are known regulators of neural stem cell proliferation^50-52^. HNs were enriched for clathrin-dependent endocytosis and IRE1-mediated unfolded protein response (**Supplemental Table 6**). Endocytosis is critical for neuronal vesicle recycling at synapses and endoplasmic reticulum stress affects the response of the hypothalamus to external stimuli in obesity^53^. Thus, pathway analysis confirms that the implicated effector genes are likely to be important for hypothalamic development and function.

### Colocalization of loci associated with multiple traits

Next, we identified contacted genes implicated in the context of multiple GWAS traits. Although most implicated genes were specific to individual traits, we identified multiple genes that were shared, suggesting a degree of overlap in the regulatory mechanisms controlling these traits **(Figure 4C; Supplemental Table 7)**. In particular, two loci contacted four genes (*BSN*/*FAM212A* and *FEZF1* /*FEZF1-AS1*) which were identified in our scans of BMI, height, AAM and sleep. To determine whether these overlaps represent likely shared regulatory regions, we performed Hypothesis Prioritization in multi-trait Colocalization (HyPrColoc) analysis for several regions. Our results highlighted both shared and distinct regulatory architectures across traits that varied by locus. For example, the *FEZF1* region colocalized among BMI, height, and AAM (regional posterior probability (PP) = 0.91), indicating a likely shared regulatory region impacting each trait (**Figure 4D**). Interestingly, the proxy of the *FEZF1* signal was located in a putative NRF1 binding site (**Figure 4D; Supplemental Table 8**). In contrast to the *FEZF1* locus, although the well-known known *BDNF* was implicated as an effector gene for AAM, BMI, and sleep, these three signals appeared to be distinct (PP = 0), suggesting a complex regulatory architecture for this region that differs by trait (**Figure 4E**).

### Colocalization of target effector genes with eQTLs – cumulative evidence

Multiple data sources can contribute orthogonal evidence for effector genes at GWAS loci^49^. The GTEx consortium^35^ has characterized hypothalamic tissue eQTLs, so we performed colocalization analyses to assess how many gene-SNP connections agreed with the physical variant-to-gene mapping approach in our specific cellular settings. For AAM, 13 genes colocalized with eQTLs, with two adjacent genes supported by our variant-to-gene mapping approach, *RPS26* (PP = 0.951) and *SUOX* (PP = 0.942). For BMI, we observed 12 colocalized genes, with one gene supported by our variant-to-gene mapping approach, *DHRS11* (PP = 0.822). Of the 29 genes colocalized with eQTLs for height, three were supported by our data: *NMT1* (PP = 0.94), *RFT1* (PP = 0.85) and *RPS9* (PP = 0.75). There was only one eQTL colocalized for sleep, but was not detected by our approach. For MDD and bipolar disorder, no genes colocalized with the eQTL data, which may be due to the relatively few signals detected in the eQTL analysis.

## DISCUSSION

We used an established *in vitro* HN model in order to both understand its genomic architecture and to gain insight into mechanisms by which non-coding GWAS loci associated with hypothalamic-regulated traits could mediate their effects. Given the challenge in acquiring primary human hypothalamic tissue and the organ’s complex makeup of cell and neuronal types^54^, we leveraged RNA-seq, ATAC-seq and promoter-focused Capture C to identify an aggregation of potentially relevant cis-regulatory regions in ESC-derived HNs. Importantly, we verified that HNs exhibited temporal transcriptional profiles that are congruent with *in vivo* hypothalamic molecular expression signatures and functional networks^27^.

We mapped common GWAS variants associated with AAM, BMI, height, bipolar disorder, sleep, and MDD to putative effector genes via their likely cREs. This approach identified both known and novel genes. For example, *FEZF1* mutations cause hypogonadotropic hypogonadism with anosmia^47^. *FEZF1* is a zinc finger transcriptional repressor that is critical for hypothalamus development^55^. The proxy contacting the *FEZF1* promoter is located in a binding site for NRF1, a transcription factor that regulates expression of several genes involved in mitochondrial biosynthesis and respiration, but is also important for neuronal differentiation and axogenesis^56^. *FEZF1* mutations impair puberty by disrupting the migration of gonadotropin-releasing hormone neurons, which are necessary to initiate puberty, from the olfactory bulb placode to the hypothalamus during fetal development^47^. In contrast, while *BDNF* was implicated in three traits, we observed distinct GWAS association landscapes, with different sentinels pointing to different proxies that consistently contacted the *BDNF* promoter. Thus, *BDNF* appears to have an intricate regulatory architecture and harbors multiple trait-associated variants that likely act in cell-type and temporally-specific contexts.

To uncover genes implicated by multiple analytic approaches, we also performed colocalization analyses of the implicated traits with hypothalamic eQTLs. Both eQTL and variant-to-gene mapping approaches identified *DHRS11* for BMI. The overlap between the two approaches was low, possibly due to differences between *ex vivo* tissue samples and stem cell-derived cells. A confluence of evidence is critical for distinguishing true effector genes from the many ‘bystander genes’ identified in eQTL studies^49^; our physical variant-to-gene mapping pipeline represents one such approach.

Here we report aspects of genomic architecture of a stem cell-based model of human hypothalamic development. We relate this architecture to the cellular ontogenesis of the human hypothalamus, and to the regulation of genes that influence complex phenotypes. Application of these strategies enables specific gene attributions for non-coding SNPs implicated in relevant common traits by GWAS efforts. These integrated datasets therefore offer valuable insight for prioritizing candidate genes that drive the molecular mechanisms by which the hypothalamus contributes to the pathogenesis of relevant complex traits.

## ACKNOWLEDGEMENTS

We acknowledge Elisabetta Manduchi for establishing the Capture C pipeline. The project described was supported by the National Center for Research Resources, Grant UL1RR024134, and is now at the National Center for Advancing Translational Sciences, Grant UL1TR000003. The content is solely the responsibility of the authors and does not necessarily represent the official views of the NIH. Supported in part by the Institute for Translational Medicine and Therapeutics’ (ITMAT) Transdisciplinary Program in Translational Medicine and Therapeutics. D.L.C. is supported by the NICHD (NIH1K99HD099330-01). R.L.L is supported by DK52431-23 and P30DK026687-41. S.F.A.G. is supported by R01-HD056465, R01 HG010067, and the Daniel B. Burke Endowed Chair for Diabetes Research.

## SUPPLEMENTAL FIGURE LEGENDS

**Supplemental Figure 1:** (A) The first and second principal components of RNA-seq and (B) ATAC-seq samples. The variance explained by each principal component is indicated on each axis. (C) Pairwise Pearson correlation coefficients of RNA-seq and (D) ATAC-seq samples. (E) The expression (TPM) of selected marker genes during the three stages of differentiation.

**Supplemental Figure 2:** (A) Counts of the number of genes assigned to each cluster. (B) The relative number of genes annotated as either protein coding, lincRNA, or other annotation (e.g. small noncoding RNAs, pseudogenes). (C) Top 10 GO term or (D) REACTOME pathway enrichment of each gene group. The bar’s length depicts the -log10 transformed FDR adjusted p-values. The red dotted line indicates the threshold for statistical significance (FDR < 0.05).

**Supplemental Figure 3:** (A) The genomic location of all OCRs and the (C) PIR-OCRs. (B) Distribution of the number of cRE at each stage. (C) Quantification of the distance (log10) of each cRE (D) Distance between fragments detected by Promoter-focused Capture C at 1-fragment and 4-fragment resolutions from each stage of differentiation. (E) Distribution of the number of PIR-OCRs linked to each gene in each stage of differentiation. (F) The distribution of the number of cRE binned by expression quantile. (G) Chromatin accessibility profile relative to all transcription start sites. (H) Percentage of fragments with interactions to other promoter bait fragments (bait-to-bait) at each resolution.

**Supplemental Figure 4:** (A) Comparison between ESC and HP enrichment scores (-log_10_(BiFET P value)). (B) Comparison between HN and HP enrichment scores. (C-F) Expression (log_2_TPM) of top enriched TFs biased toward HPs compared to ESC (C), HP compared to HN (D) and HN to HP (E), and TFs highly enriched in both HPs and HNs (F).

## SUPPLEMENTARY TABLES

**Supplemental Table 1:** Results from Spearman Correlation tests between HNs and GTeX tissues.

**Supplemental Table 2:** Enriched TF motifs in cRE compared to unannotated OCRs

**Supplemental Table 3:** Results for partitioned LD score regression for the indicated genotypes.

**Supplemental Table 4:** DAVID annotations of implicated genes for each trait.

**Supplemental Table 5:** Gene annotation to each indicated proxy and sentinel SNP.

**Supplemental Table 6:** Overlap between GWAS traits, 0 = not implicated, 1 = implicated.

**Supplemental Table 7:** Annotation of SNPs located in TF binding sites.

**Supplemental Table 8:** Enriched GO terms of genes implicated by our V2G pipeline mapping specifically in HPs and HNs.

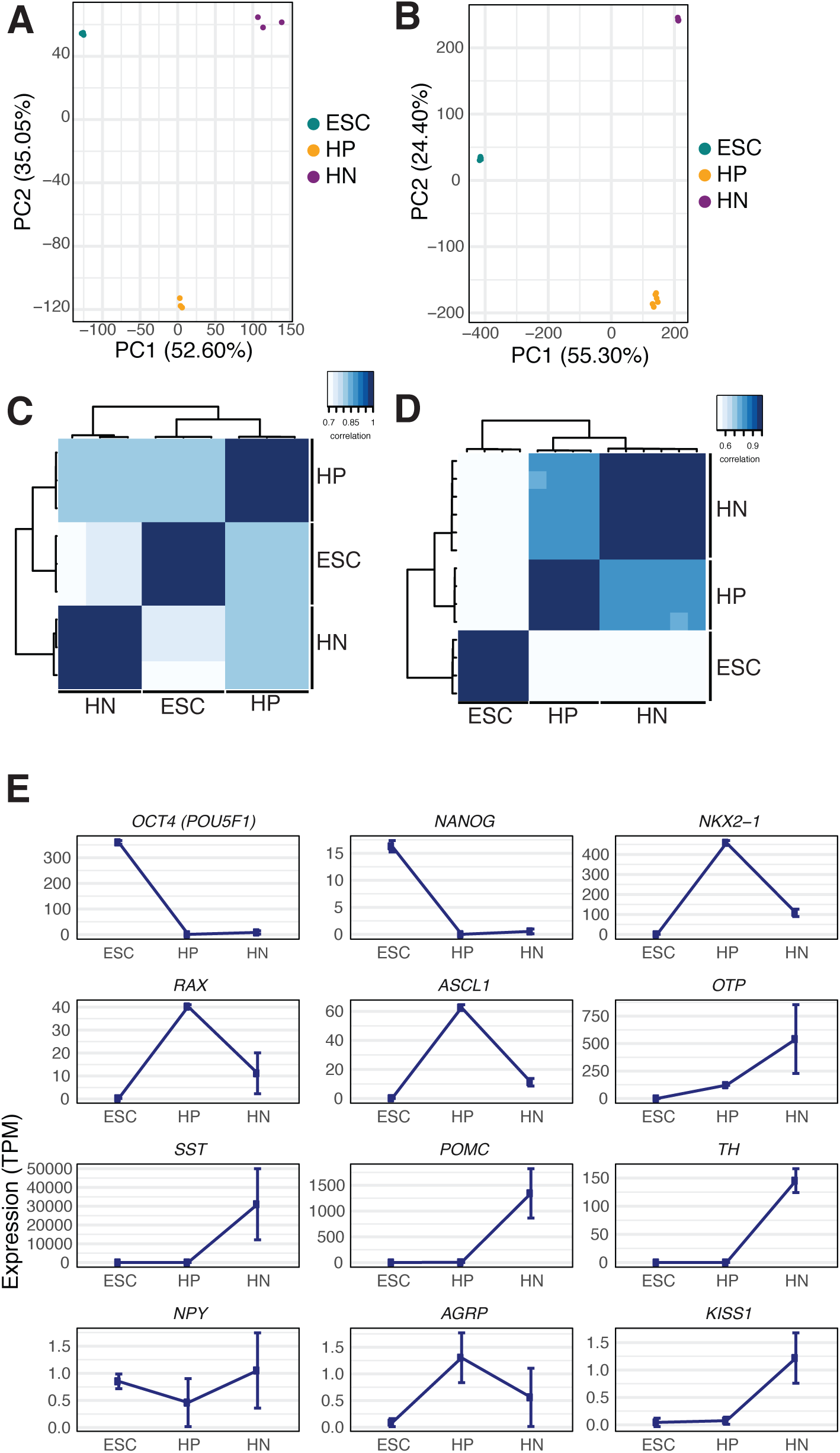

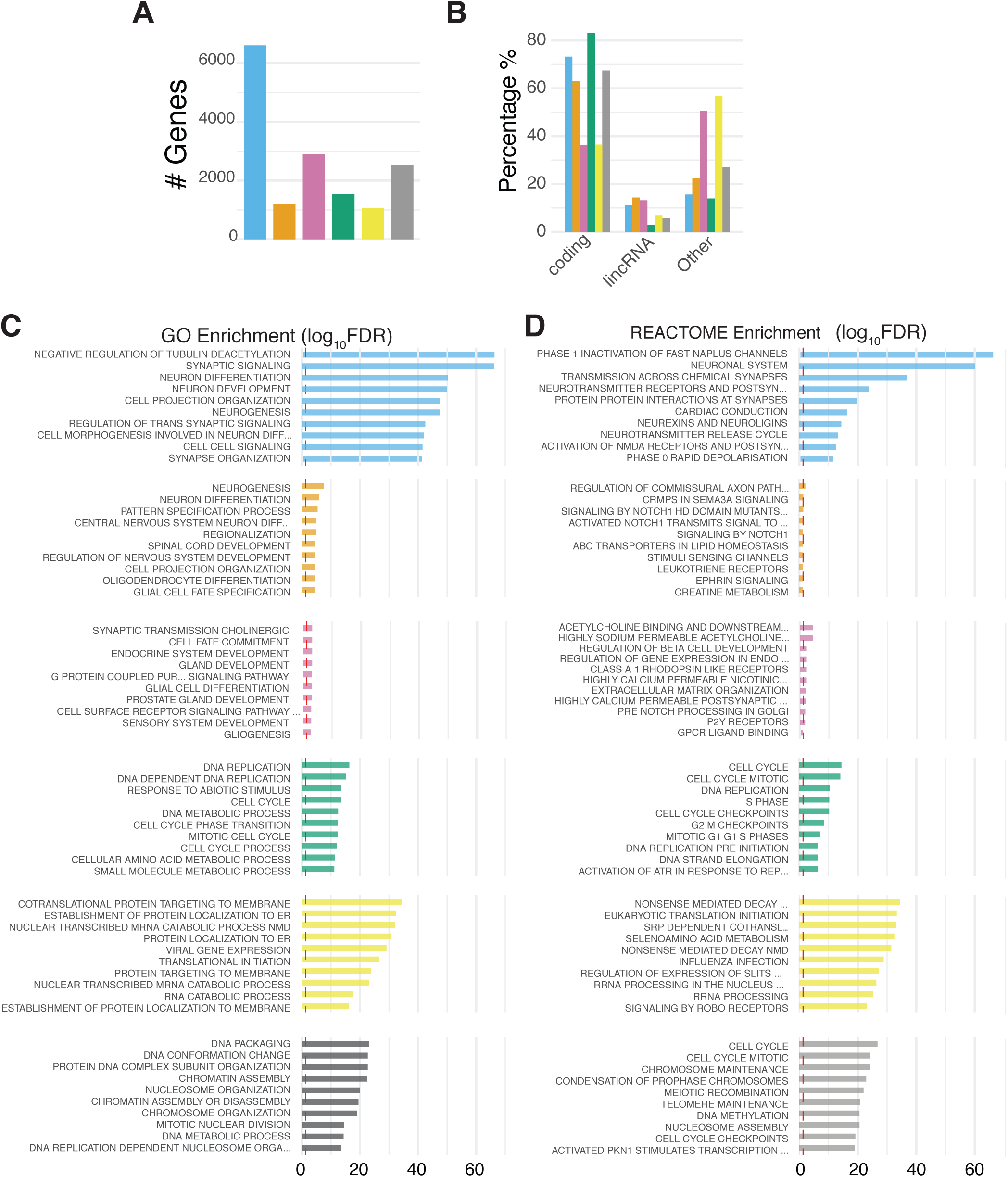

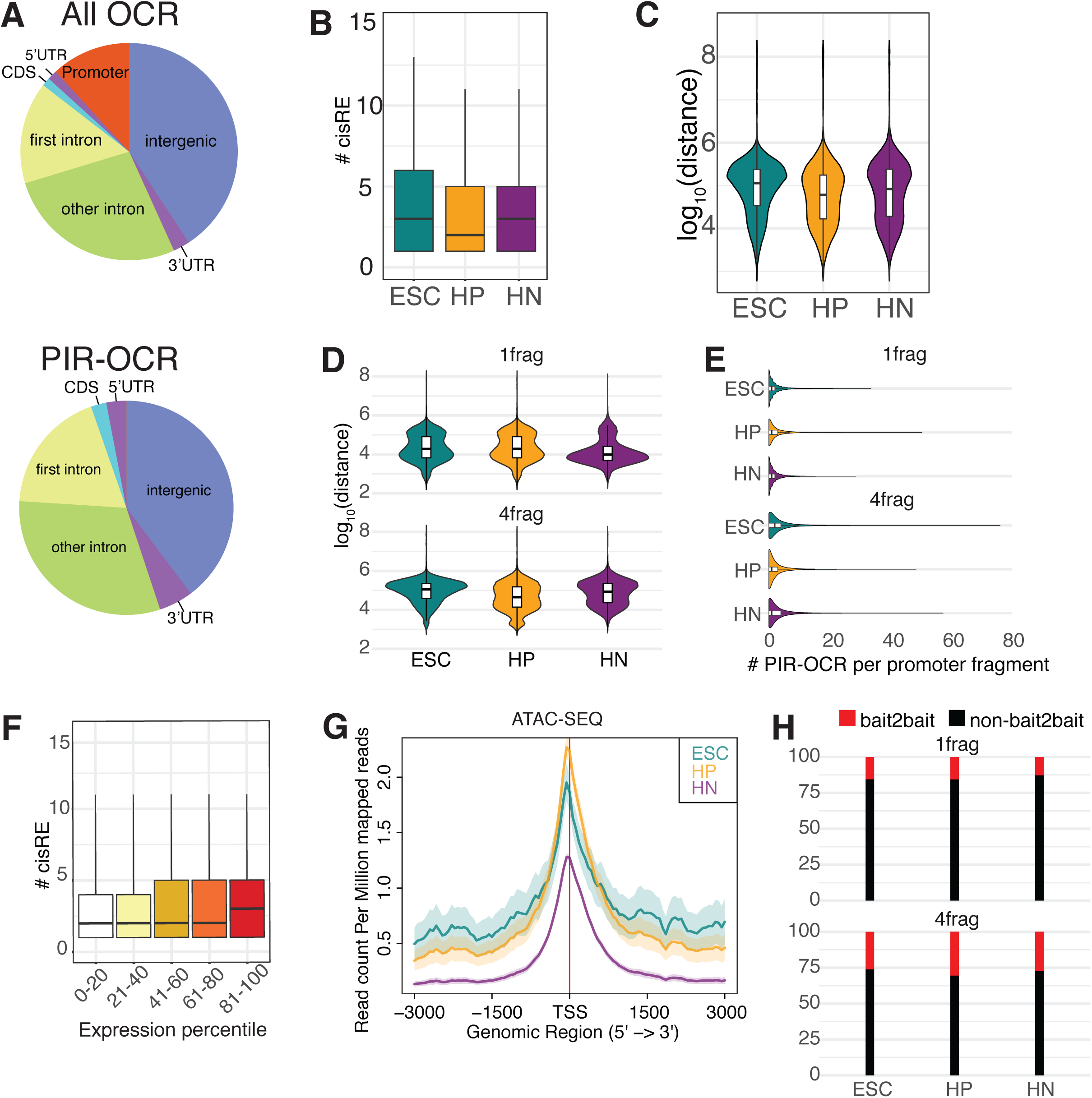

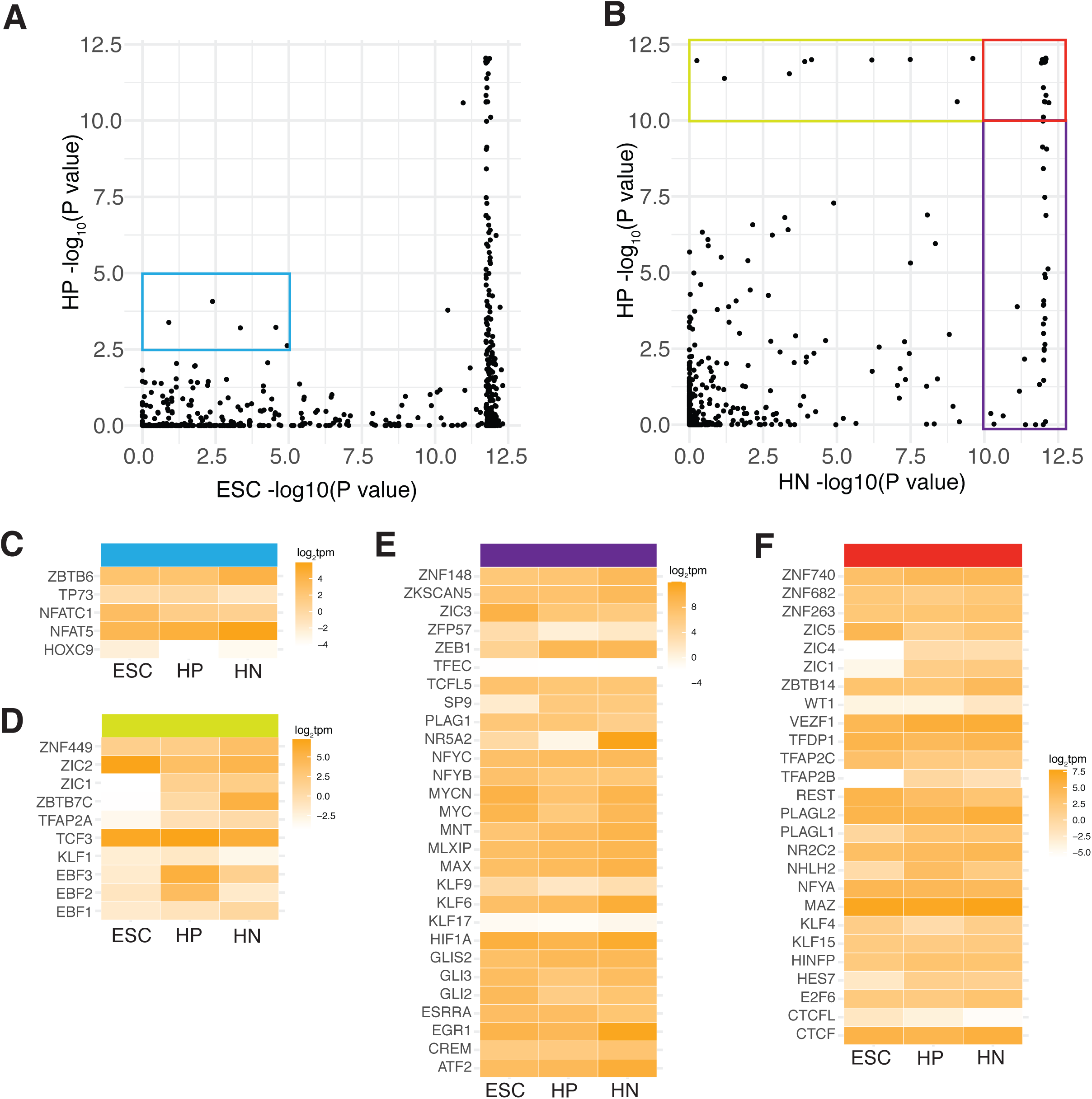

## REFERENCES

1. Merkle FT, Maroof A, Wataya T, Sasai Y, Studer L, Eggan K et al. Generation of neuropeptidergic hypothalamic neurons from human pluripotent stem cells. Development 2015; 142(4): 633–643.

2. Andermann ML, Lowell BB. Toward a Wiring Diagram Understanding of Appetite Control. Neuron 2017; 95(4): 757–778.

3. Yoo S, Blackshaw S. Regulation and function of neurogenesis in the adult mammalian hypothalamus. Prog Neurobiol 2018; 170: 53–66.

4. Herbison AE. Control of puberty onset and fertility by gonadotropin-releasing hormone neurons. Nat Rev Endocrinol 2016; 12(8): 452–466.

5. Rajamani U, Gross AR, Hjelm BE, Sequeira A, Vawter MP, Tang J et al. Super-Obese Patient-Derived iPSC Hypothalamic Neurons Exhibit Obesogenic Signatures and Hormone Responses. Cell Stem Cell 2018; 22(5): 698-712.e699.

6. Rajamani U, Gross AR, Hjelm BE, Sequeira A, Vawter MP, Tang J et al. Super-Obese Patient-Derived iPSC Hypothalamic Neurons Exhibit Obesogenic Signatures and Hormone Responses. Cell Stem Cell 2018; 22(5): 698–712 e699.

7. Wang L, Egli D, Leibel RL. Efficient Generation of Hypothalamic Neurons from Human Pluripotent Stem Cells. Curr Protoc Hum Genet 2016; 90: 21 25 21–21 25 14.

8. Dashti HS, Jones SE, Wood AR, Lane JM, van Hees VT, Wang H et al. Genome-wide association study identifies genetic loci for self-reported habitual sleep duration supported by accelerometer-derived estimates. Nat Commun 2019; 10(1): 1100.

9. Jones SE, Lane JM, Wood AR, van Hees VT, Tyrrell J, Beaumont RN et al. Genome- wide association analyses of chronotype in 697,828 individuals provides insights into circadian rhythms. Nat Commun 2019; 10(1): 343.

10. Jones SE, van Hees VT, Mazzotti DR, Marques-Vidal P, Sabia S, van der Spek A et al. Genetic studies of accelerometer-based sleep measures yield new insights into human sleep behaviour. Nat Commun 2019; 10(1): 1585.

11. Wang L, De Solis AJ, Goffer Y, Birkenbach KE, Engle SE, Tanis R et al. Ciliary gene RPGRIP1L is required for hypothalamic arcuate neuron development. JCI Insight 2019; 4(3).

12. Stratigopoulos G, Burnett LC, Rausch R, Gill R, Penn DB, Skowronski AA et al. Hypomorphism of Fto and Rpgrip1l causes obesity in mice. J Clin Invest 2016; 126(5): 1897–1910.

13. Claussnitzer M, Dankel SN, Kim KH, Quon G, Meuleman W, Haugen C et al. FTO Obesity Variant Circuitry and Adipocyte Browning in Humans. N Engl J Med 2015; 373(10): 895–907.

14. Smemo S, Tena JJ, Kim KH, Gamazon ER, Sakabe NJ, Gomez-Marin C et al. Obesity- associated variants within FTO form long-range functional connections with IRX3. Nature 2014; 507(7492): 371–375.

15. Frayling TM, Timpson NJ, Weedon MN, Zeggini E, Freathy RM, Lindgren CM et al. A common variant in the FTO gene is associated with body mass index and predisposes to childhood and adult obesity. Science 2007; 316(5826): 889–894.

16. Siersbaek R, Madsen JGS, Javierre BM, Nielsen R, Bagge EK, Cairns J et al. Dynamic Rewiring of Promoter-Anchored Chromatin Loops during Adipocyte Differentiation. Mol Cell 2017; 66(3): 420–435 e425.

17. Javierre BM, Burren OS, Wilder SP, Kreuzhuber R, Hill SM, Sewitz S et al. Lineage- Specific Genome Architecture Links Enhancers and Non-coding Disease Variants to Target Gene Promoters. Cell 2016; 167(5): 1369–1384 e1319.

18. Schmitt AD, Hu M, Jung I, Xu Z, Qiu Y, Tan CL et al. A Compendium of Chromatin Contact Maps Reveals Spatially Active Regions in the Human Genome. Cell Rep 2016; 17(8): 2042–2059.

19. Chesi A, Wagley Y, Johnson ME, Manduchi E, Su C, Lu S et al. Genome-scale Capture C promoter interactions implicate effector genes at GWAS loci for bone mineral density. Nat Commun 2019; 10(1): 1260.

20. Caliskan M, Manduchi E, Rao HS, Segert JA, Beltrame MH, Trizzino M et al. Genetic and Epigenetic Fine Mapping of Complex Trait Associated Loci in the Human Liver. Am J Hum Genet 2019; 105(1): 89–107.

21. Cousminer DL, Wagley Y, Pippin JA, Elhakeem A, Way GP, McCormack SE et al. Genome-wide association study implicates novel loci and reveals candidate effector genes for longitudinal pediatric bone accrual through variant-to-gene mapping. medRxiv 2020.

22. Su C, Johnson ME, Torres A, Thomas RM, Manduchi E, Sharma P et al. Human follicular helper T cell promoter connectomes reveal novel genes and regulatory elements at SLE GWAS loci. bioRxiv 2019.

23. Jung I, Schmitt A, Diao Y, Lee AJ, Liu T, Yang D et al. A compendium of promoter- centered long-range chromatin interactions in the human genome. Nat Genet 2019; 51(10): 1442–1449.

24. Bonev B, Mendelson Cohen N, Szabo Q, Fritsch L, Papadopoulos GL, Lubling Y et al. Multiscale 3D Genome Rewiring during Mouse Neural Development. Cell 2017; 171(3): 557-572.e524.

25. Shimogori T, Lee DA, Miranda-Angulo A, Yang Y, Wang H, Jiang L et al. A genomic atlas of mouse hypothalamic development. Nat Neurosci 2010; 13(6): 767–775.

26. Huisman C, Cho H, Brock O, Lim SJ, Youn SM, Park Y et al. Single cell transcriptome analysis of developing arcuate nucleus neurons uncovers their key developmental regulators. Nat Commun 2019; 10(1): 3696.

27. Wang L, Meece K, Williams DJ, Lo KA, Zimmer M, Heinrich G et al. Differentiation of hypothalamic-like neurons from human pluripotent stem cells. J Clin Invest 2015; 125(2): 796–808.

28. Sherwood RI, Hashimoto T, O’Donnell CW, Lewis S, Barkal AA, van Hoff JP et al. Discovery of directional and nondirectional pioneer transcription factors by modeling DNase profile magnitude and shape. Nat Biotechnol 2014; 32(2): 171–178.

29. Fornes O, Castro-Mondragon JA, Khan A, van der Lee R, Zhang X, Richmond PA et al. JASPAR 2020: update of the open-access database of transcription factor binding profiles. Nucleic Acids Res 2020; 48(D1): D87–D92.

30. Finucane HK, Bulik-Sullivan B, Gusev A, Trynka G, Reshef Y, Loh PR et al. Partitioning heritability by functional annotation using genome-wide association summary statistics. Nat Genet 2015; 47(11): 1228–1235.

31. Giambartolomei C, Vukcevic D, Schadt EE, Franke L, Hingorani AD, Wallace C et al. Bayesian test for colocalisation between pairs of genetic association studies using summary statistics. PLoS Genet 2014; 10(5): e1004383.

32. Burnett LC, Leduc CA, Sulsona CR, Paull D, Rausch R, Eddiry S et al. Deficiency in prohormone convertase PC1 impairs prohormone processing in Prader-Willi syndrome. Journal of Clinical Investigation 2016; 127(1): 293–305.

33. Burnett LC, LeDuc CA, Sulsona CR, Paull D, Rausch R, Eddiry S et al. Deficiency in prohormone convertase PC1 impairs prohormone processing in Prader-Willi syndrome. J Clin Invest 2017; 127(1): 293–305.

34. Yee CL, Wang Y, Anderson S, Ekker M, Rubenstein JL. Arcuate nucleus expression of NKX2.1 and DLX and lineages expressing these transcription factors in neuropeptide Y(+), proopiomelanocortin(+), and tyrosine hydroxylase(+) neurons in neonatal and adult mice. J Comp Neurol 2009; 517(1): 37–50.

35. Consortium GT, Laboratory DA, Coordinating Center -Analysis Working G, Statistical Methods groups-Analysis Working G, Enhancing Gg, Fund NIHC et al. Genetic effects on gene expression across human tissues. Nature 2017; 550(7675): 204–213.

36. Subramanian A, Tamayo P, Mootha VK, Mukherjee S, Ebert BL, Gillette MA et al. Gene set enrichment analysis: a knowledge-based approach for interpreting genome-wide expression profiles. Proc Natl Acad Sci U S A 2005; 102(43): 15545–15550.

37. Park SG, Hannenhalli S, Choi SS. Conservation in first introns is positively associated with the number of exons within genes and the presence of regulatory epigenetic signals. BMC Genomics 2014; 15: 526.

38. Song M, Yang X, Ren X, Maliskova L, Li B, Jones IR et al. Mapping cis-regulatory chromatin contacts in neural cells links neuropsychiatric disorder risk variants to target genes. Nat Genet 2019; 51(8): 1252–1262.

39. Ma Y, McKay DJ, Buttitta L. Changes in chromatin accessibility ensure robust cell cycle exit in terminally differentiated cells. PLoS Biol 2019; 17(9): e3000378.

40. Slattery M, Zhou T, Yang L, Dantas Machado AC, Gordan R, Rohs R. Absence of a simple code: how transcription factors read the genome. Trends Biochem Sci 2014; 39(9): 381–399.

41. Briscoe J, Pierani A, Jessell TM, Ericson J. A Homeodomain Protein Code Specifies Progenitor Cell Identity and Neuronal Fate in the Ventral Neural Tube. Cell 2000; 101(4): 435–445.

42. Hilger-Eversheim K. MM, Schorle H., Buettner R. Regulatory roles of AP-2 transcription factors in vertebrate development, apoptosis, and cell cycle control. Genes Cells 2000; 260: 1–12.

43. Luo Z, Gao X, Lin C, Smith ER, Marshall SA, Swanson SK et al. Zic2 is an enhancer- binding factor required for embryonic stem cell specification. Mol Cell 2015; 57(4): 685–694.

44. Leslie J. Baier YLM, Maria Sara Remedi, Michael Traurig, Paolo Piaggi GW, Ke Huang, Alyssa Stacy, Sayuko Kobes, Jonathan Krakoff PHB, Robert G. Nelson, William C. Knowler, Robert L. Hanson CGN, and Clifton Bogardus. ABCC8 R1420H Loss-of- Function Variant in a Southwest American Indian Community: Association With Increased Birth Weight and Doubled Risk of Type 2 Diabetes. Diabetes 2015; 64: 4322–4332.

45. Sandrini L, Di Minno A, Amadio P, Ieraci A, Tremoli E, Barbieri SS. Association between Obesity and Circulating Brain-Derived Neurotrophic Factor (BDNF) Levels: Systematic Review of Literature and Meta-Analysis. Int J Mol Sci 2018; 19(8).

46. Moller DE, Berger JP. Role of PPARs in the regulation of obesity-related insulin sensitivity and inflammation. Int J Obes Relat Metab Disord 2003; 27 Suppl 3: S17–21.

47. Kotan LD, Hutchins BI, Ozkan Y, Demirel F, Stoner H, Cheng PJ et al. Mutations in FEZF1 cause Kallmann syndrome. Am J Hum Genet 2014; 95(3): 326–331.

48. Miyagawa T, Hida A, Shimada M, Uehara C, Nishino Y, Kadotani H et al. A missense variant in PER2 is associated with delayed sleep-wake phase disorder in a Japanese population. J Hum Genet 2019; 64(12): 1219–1225.

49. Ndungu A, Payne A, Torres JM, van de Bunt M, McCarthy MI. A Multi-tissue Transcriptome Analysis of Human Metabolites Guides Interpretability of Associations Based on Multi-SNP Models for Gene Expression. Am J Hum Genet 2020; 106(2): 188–201.

50. Sato A, Sunayama J, Matsuda K, Tachibana K, Sakurada K, Tomiyama A et al. Regulation of neural stem/progenitor cell maintenance by PI3K and mTOR. Neurosci Lett 2010; 470(2): 115–120.

51. Imamura O, Pages G, Pouyssegur J, Endo S, Takishima K. ERK1 and ERK2 are required for radial glial maintenance and cortical lamination. Genes Cells 2010; 15(10): 1072–1088.

52. Le Belle JE, Orozco NM, Paucar AA, Saxe JP, Mottahedeh J, Pyle AD et al. Proliferative neural stem cells have high endogenous ROS levels that regulate self-renewal and neurogenesis in a PI3K/Akt-dependant manner. Cell Stem Cell 2011; 8(1): 59–71.

53. Ozcan L, Ergin AS, Lu A, Chung J, Sarkar S, Nie D et al. Endoplasmic reticulum stress plays a central role in development of leptin resistance. Cell Metab 2009; 9(1): 35–51.

54. Campbell JN, Macosko EZ, Fenselau H, Pers TH, Lyubetskaya A, Tenen D et al. A molecular census of arcuate hypothalamus and median eminence cell types. Nat Neurosci 2017; 20(3): 484–496.

55. Hirata T, Nakazawa M, Muraoka O, Nakayama R, Suda Y, Hibi M. Zinc-finger genes Fez and Fez-like function in the establishment of diencephalon subdivisions. Development 2006; 133(20): 3993–4004.

56. Kiyama T, Chen CK, Wang SW, Pan P, Ju Z, Wang J et al. Essential roles of mitochondrial biogenesis regulator Nrf1 in retinal development and homeostasis. Mol Neurodegener 2018; 13(1): 56.

